# Brain dynamics supporting high cognitive performance reorganize after midlife

**DOI:** 10.64898/2026.06.02.729404

**Authors:** Yaroslav Chekin, Dakota Decker, Julien Dubois, Ryan M. Field, Andrew Gundran, Austin Jewison, Erin M. Koch, Gabriel Lerner, Zahra M. Aghajan, Naomi Miller, Katherine L. Perdue, Moriah Taylor

**Affiliations:** Kernel, 10361 Jefferson Blvd, Culver City, CA 90232 USA

## Abstract

Quantifying functional brain aging trajectories at scale remains a fundamental challenge due to the scanner-bound limitations of traditional neuroimaging. Here, we deploy whole-head Time-Domain functional Near-Infrared Spectroscopy (TD-fNIRS) to map task-evoked cortical dynamics during a 30-minute cognitive battery across the adult lifespan (N = 302, age 18–87, 45% racial or ethnic minority). We developed a robust General Cognitive Factor (GCF) tracking age-related performance decline (r = −0.57, p < 0.0001). Analysis of brain activity patterns revealed systemic, age-dependent neural dedifferentiation, highlighting a distinct neurocognitive inflection point around age 55. Prior to this threshold, high performers exhibit more variable neural activation across tasks; post-age 55, high performance is sustained through reduced spatial differentiation, signaling a compensatory strategy. Furthermore, subjective anxiety and depression disrupt these compensatory mechanisms, and subjective cognitive complaints register as distinct neural signatures before behavioral GCF decline manifests. Together, these findings establish a scalable framework for mapping functional brain health across the human lifespan, uncovering the neural mechanics of cognitive resilience and vulnerability before behavioral decline manifests.

## Introduction

Healthy aging is known to change cognitive performance and the underlying patterns of neural activity in complex ways. Large datasets of structural magnetic resonance images have shown broad anatomical changes^1^, and resting state functional magnetic resonance imaging (fMRI) has shown shifts in connectivity with aging ^2–5^. However, neither of these methods is sufficient to capture the cognitive effects most relevant to the experience of aging humans in their everyday lives. To probe these neural changes directly, it is imperative to measure the brain while it is performing cognitive tasks^2^. Direct access to the activity of the brain during these tasks can also shed light on theoretical models of cognitive aging, as different patterns of brain activity may underlie similar behavior^6,7^.

Despite the promise of the approach, task-based brain measurement has proven to be challenging due to the high expense of scans and the necessity of carefully designing tasks to capture a range of cognitive functions. While the existing large scale studies such as the UK Biobank^4^ and the Cambridge Centre for Ageing and Neuroscience (Cam-CAN)^8,9^ provide a wealth of data for scientific discovery, it is unlikely that fMRI could be used routinely at scale to monitor cognition in the aging population due to cost and accessibility constraints. Cognitive screening currently relies on coarse behavioral screening tools such as the Mini Mental State Exam^10^ and Montreal Cognitive Assessment^11^, and does not capture subtle changes in cognition. Given that changes in brain functional responses have been shown to precede cognitive decline, broadening access to these measures could allow for interventions to preserve cognition^12^.

One scalable approach to measuring brain activity in a broad population is using functional near-infrared spectroscopy (fNIRS). The lower cost of the technology and ease of setup not only increases potential for widespread use, but also enables collection of large datasets that are well-suited for interrogating complex changes in the brain with aging^13^. fNIRS has been used extensively to study both healthy aging^14^ and cognitive impairment^15–17^. Direct measurement of both oxygenated and deoxygenated hemoglobin not only captures changes in functional brain activity, but also means the technology is sensitive to vascular dynamics, which have been shown to causally precede cognitive decline^18^. Yet, as a whole, existing fNIRS studies typically have limitations in the number of brain regions measured, as most systems do not provide full-head coverage, and limitations in scope to single tasks or cognitive domain ^14,19^.

Several theories of cognitive ability and healthy aging involve interplay between multiple cognitive domains, such as changes in attention leading to poor memory performance^9,20^. Established theories of the aging brain typically involve large-scale shifts in brain recruitment, such as the hemispheric asymmetry reduction in older adults (HAROLD), compensation-related utilization of neural circuits hypothesis (CRUNCH), the Scaffolding Theory of Aging and Cognition (STAC), and more generally dedifferentiation, the blurring of specialized brain systems^5,21–23^. However, others have found evidence for the maintenance hypothesis, which claims that retention of dynamic responsivity in specialized networks is key to successful cognitive aging^24,25^. Therefore, capturing the full nature of cognitive aging requires measurement of a broad range of cognitive skills alongside the whole-brain activation underlying their execution.

To address this gap, we collected a large dataset of healthy adults (N=302, age 18-87, 45% participants from racial or ethnic minority groups) completing a 30-minute multifactor cognitive battery. Brain activity was measured simultaneously with a portable time domain (TD)-fNIRS system, allowing for high-fidelity measurements over the whole cortex^26^. A subset of participants (n=106) were measured twice, approximately 1 month apart to quantify reliability. We first present a general behavioral cognitive factor that summarizes the overall performance across domains, along with a general brain factor that characterizes differentiation of task-evoked brain activity across the range of cognitive tasks. We examined how both factors changed across the lifespan, the complex interactions between these factors and age, and whether these factors are sensitive to subjective reports of affect and cognition. Finally, we present evidence that together these general factors can be used to summarize and track cognitive aging in the adult population or in response to interventions that may be expected to improve cognition. Overall, this advance represents an important step forward in routine monitoring of brain health with aging.

## Results

Healthy adult participants (n=302) performed a streamlined battery of cognitive tasks (e.g. language production, cognitive control, memory, and fluid intelligence; Methods, Fig 1), on the computer while their brain activity was simultaneously recorded via TD-fNIRS. Participants were intentionally selected to cover the range of adult lifespan, 18 years of age to 87 years of age (mean 48.2; Fig. 1b, left) and recruitment attempts were made to balance biological sex within each decade of life (Fig. 1b, right) shown here over decades. Task instructions, task practices, and intra-task breaks were automated and self-paced, minimizing the role of the operator in the data acquisition process. Additional demographic information including highest level of education achieved and ethnicity and race were also collected via short self-administered surveys, alongside a series of clinically validated surveys that captured broader markers of well-being, including the Patient Health Questionnaire (PHQ-9), Generalized Anxiety Disorder scale (GAD-7), and the Subjective Cognitive Decline Questionnaire (SCDQ-9) (Table 1). Participants across all ages were able to successfully complete the cognitive battery while wearing the headset.

**Figure 1:**
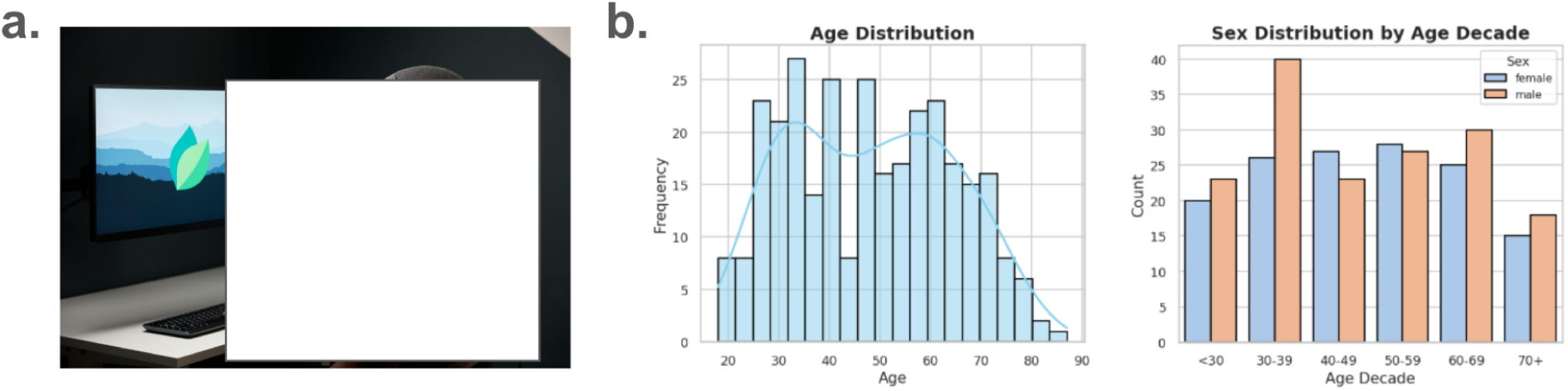
Data collection and participant ages. **a.** Model of participant wearing the headset and completing the cognitive battery. **b**. Age distribution of participants, also broken down by sex.

**Table 1:**
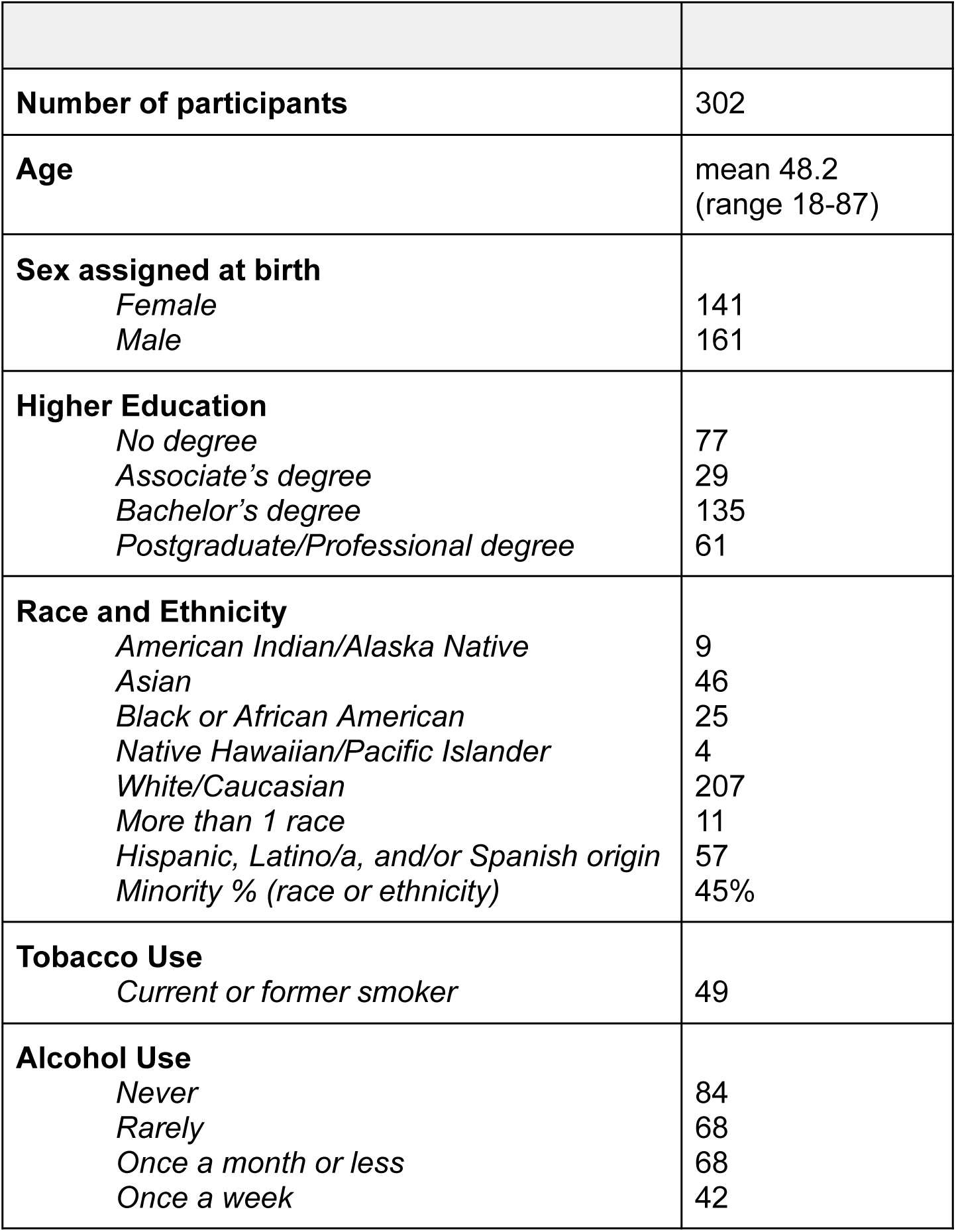
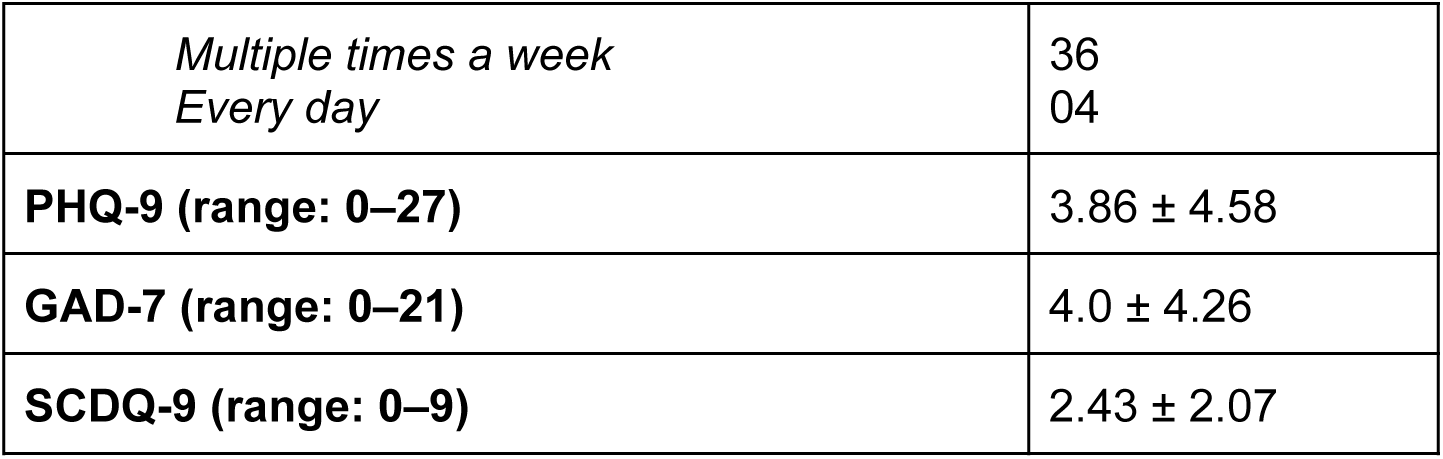
Participant demographics and characteristics.

|  |  |
| --- | --- |
| **Number of participants** | 302 |
| **Age** | mean 48.2<br>(range 18-87) |
| **Sex assigned at birth**<br>Female<br>Male | 141<br>161 |
| **Higher Education**<br>No degree<br>Associate's degree<br>Bachelor's degree<br>Postgraduate/Professional degree | 77<br>29<br>135<br>61 |
| **Race and Ethnicity**<br>American Indian/Alaska Native<br>Asian<br>Black or African American<br>Native Hawaiian/Pacific Islander<br>White/Caucasian<br>More than 1 race<br>Hispanic, Latino/a, and/or Spanish origin<br>Minority % (race or ethnicity) | 9<br>46<br>25<br>4<br>207<br>11<br>57<br>45% |
| **Tobacco Use**<br>Current or former smoker | 49 |
| **Alcohol Use**<br>Never<br>Rarely<br>Once a month or less<br>Once a week | 84<br>68<br>68<br>42 |
| *Multiple times a week*<br>*Every day* | 36<br>04 |
| **PHQ-9 (range: 0–27)** | 3.86 ± 4.58 |
| **GAD-7 (range: 0–21)** | 4.0 ± 4.26 |
| **SCDQ-9 (range: 0–9)** | 2.43 ± 2.07 |

In order to validate the implementation of the short battery of cognitive tasks, we quantified behavior in each task condition using standard metrics (when applicable), examples include accuracy, reaction time, efficiency, d-prime, word rate, inter-word variability, and search time, among many others. We then applied principal component analysis (PCA) to the set of metrics within each task condition to reduce behavioral metrics to 1–3 principal components (PCs) per task condition. Inspection of these PCs yielded interpretable factors across all task conditions. For example, the Go/No-go task was reduced to 2 PCs, capturing 1) efficacy of timing and hitting ‘go-trials’ and 2) cognitive control and the ability to inhibit responses to ‘no-go-trials’ (Fig 2a). The 2-back task was reduced to three PCs (Fig 2b): PC1 captured overall discriminability, with accuracy, d-prime, hit rate, and correct rejection rate all loading in the same direction; PC2 captured processing speed and efficiency; and PC3 captured response bias, with hit rate and correct rejection rate loading in opposite directions. In total, we reduced behavior to 22 specific task condition dimensions, which were entered into a second-level PCA to obtain a General Cognitive Factor, GCF (Fig 2c). The GCF loaded from all task conditions (though lower weights were given to verbal fluency PCs, possibly reflecting the known resilience of language in healthy aging) and was strongly correlated with age (r = −0.57, p < 0.0001; Fig 2d, left). Importantly, lower GCF values indicate poorer cognitive performance, and thus a decrease with healthy aging is expected. A subset of participants (n = 106) returned for a second session with new versions of the same tasks. The GCF showed excellent test-retest reliability (r = 0.88, p < 0.0001; ICC2 = 0.834), with a slight improvement at Visit 2 consistent with modest learning effects (Fig 2d, right).

**Figure 2:**
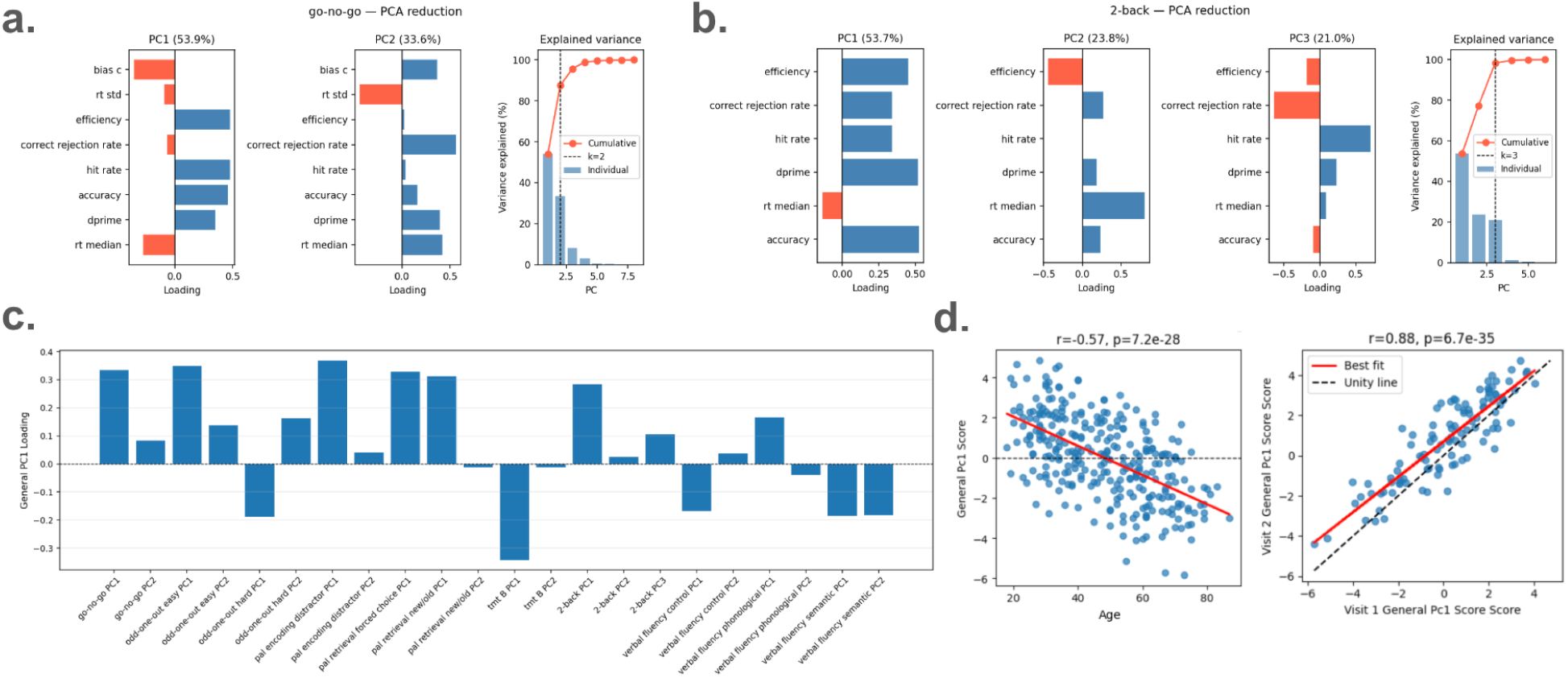
Quantification of specific task behavior and general cognition. **a.** Quantification of Go/No-go task behavior. Left: metric weights for PC1. Middle: metric weights for PC2. Right: Scree plot. **b.** Quantification of 2-back task behavior. Left: metric weights for PC1. Middle: metric weights for PC2, PC3. Right: Scree plot. In **a,b** rt stands for reaction time and bias c represents signal detection theory bias criterion. **c.** Contributions of specific task conditions to the general cognitive factor. **d.** Left: Relationship between age and the general cognitive factor. A higher general cognitive score is associated with better task performance and is negatively correlated with age. Right: Visit 1 scores versus Visit 2 scores demonstrate reliable test-retest performance (ICC2 = 0.834).

Just as performance on cognitive tasks changes with healthy aging, so too does task-related brain activation (both in terms of spatial patterns and amplitude), and in some cases neural changes occur in the absence of, or preceding, behavioral changes. In order to quantify changes in functional brain activity (relative changes in oxygenated hemoglobin; HbO) throughout the adult lifespan we executed a group level GLM for each task condition within each decade (e.g., <30, 30s, 40s, 50s, 60s, >70) and visualized the test statistics over the head.

Each task condition exhibited gradual changes in task activation with age that were generally consistent with increased activation amplitude and recruitment of additional brain regions. In the Go/No-go task (Fig. 3a), younger participants exhibited the expected right-dominant prefrontal and fronto-parietal activation that increases both in spatial extent and amplitude in the 40s and 50s, becoming widespread and bilateral by the 60s. Notably, in the oldest age group, activation has shifted entirely to non-prefrontal regions. In the ‘Hard’ Odd-One-Out task (Fig. 3b), strong activations are already present in younger age groups, likely due to the inherent difficulty of the task, with more modest recruitment of fronto-parietal and occipital regions with age.

**Figure 3:**
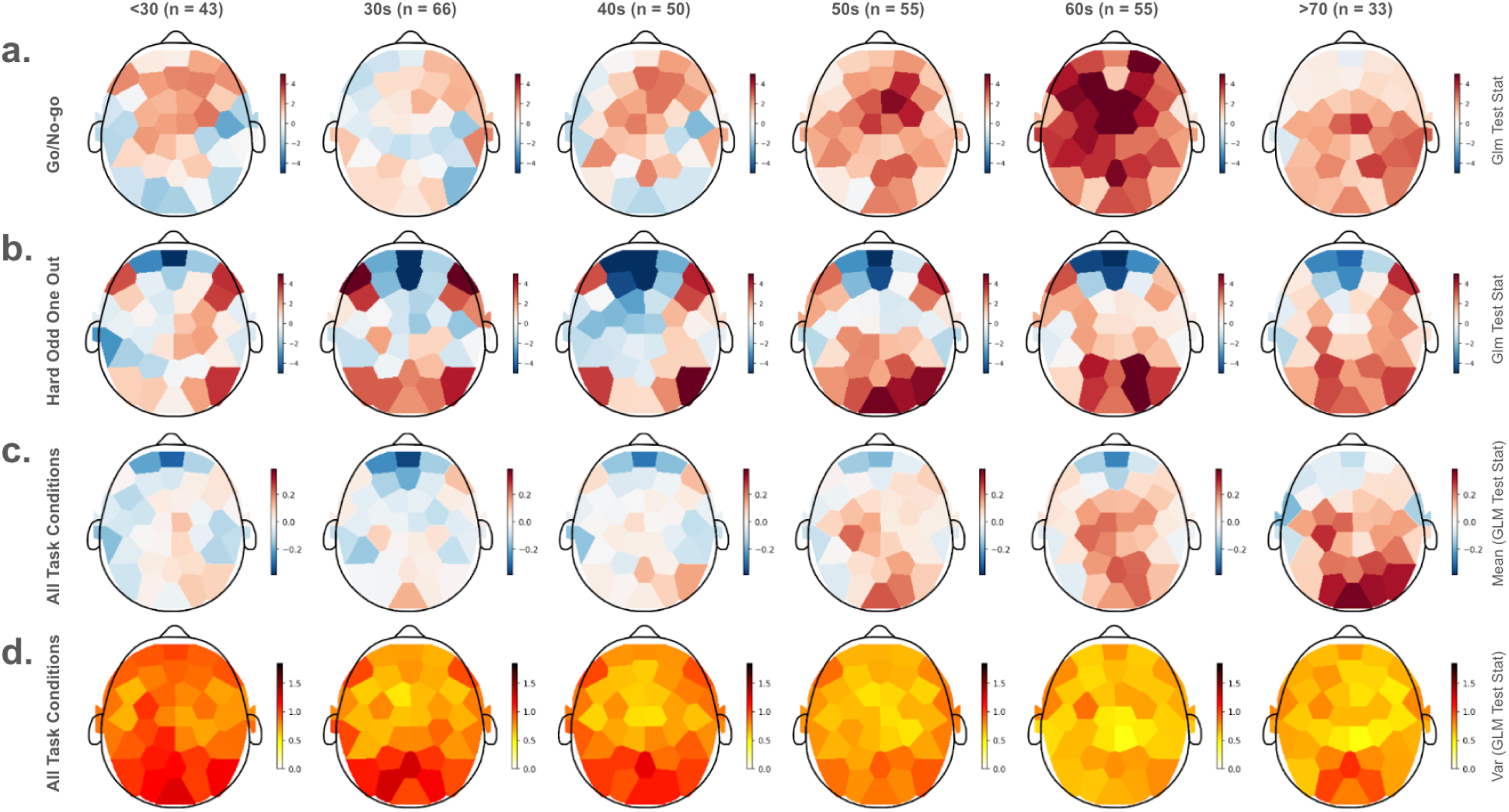
Age-related changes in task-evoked brain activity over the head. **a.** Group-level GLM statistics for HbO during the Go/No-go task, shown by decade. **b.** Same as (a) for the ‘Hard’ Odd-One-Out task condition. **c.** Mean task-evoked activity across all task conditions in the cognitive battery (average of individual task condition GLM test statistics at each channel). In **a-c** warmer colors indicate activation and cooler colors indicate deactivation. **d**. Spatial variability of task-evoked activity across all task conditions (group-average variance of individual task condition GLM statistics at each channel), shown by decade. Deeper red indicates greater average variability across tasks at that location; brighter yellow indicates more consistent activity across tasks.

The distinct task-evoked patterns of brain activity across these two tasks, apparent in the younger decades, decay with age. This visual observation of dedifferentiation of brain regions with age appeared consistent across many tasks. A simple way to quantify and visualize this is to average the patterns of brain activity across all task conditions. If patterns are highly distinct, they would largely cancel out, yielding little structure in the average. If patterns are similar across tasks, a consistent pattern would emerge. Indeed, outside of strong deactivation in medial prefrontal cortex, which likely reflects attention as this region coincides with a default mode network node, there is little-to-no general pattern of task activation in the average measurements from the younger decades (Fig. 3c, left). In contrast, beginning in middle age and progressing through the older decades, a clear general pattern of fronto-parietal and occipitoparietal activation emerges in the averages (Fig. 3c, right). As a complementary measure, we computed the variance of activity across the cognitive battery over the head for each individual, then averaged the task-evoked variability over the head within each decade. As expected, variability was greater in younger decades and, beginning in fronto-parietal regions, gradually decreased with age (Fig. 3d). Taken together, these results suggest dedifferentiation of brain activity patterns with increasing age at the group level.

Next we sought to assess individual differences in task-evoked patterns of brain activity. For each participant, we quantified pairwise distances between task-evoked activation patterns in a space defined by GLM test statistics across 35 brain regions, using two complementary distance metrics: Euclidean distance and Scaled Euclidean distance. Euclidean distance captures both magnitude and spatial configuration of activation patterns across tasks, while Scaled-Euclidean normalizes for baseline shifts in activation magnitude, thereby isolating spatial activation pattern differences independently of amplitude. We averaged these distances across all task pairs to derive a general neural differentiability score per participant and quantified its relationship with age. Consistent with the previously observed neural dedifferentiation, both metrics showed significant negative relationships with age (Euclidean: r=-0.278, p<0.001; Scaled Euclidean: r=-0.187, p=0.001), indicating that brain activity becomes less distinct across tasks with increasing age, both in terms of flexible magnitude modulation and spatial activation patterns.

Since both the general cognitive factor and the general neural differentiability scores were significantly related to age, we aimed to tease apart the interactions between neural change, cognitive performance, and healthy aging. We age-residualized the GCF and used the residuals to split participants into high performers (top 50% relative to age peers) and low performers (bottom 50% relative to age peers) and visualized group trajectories across the full age span using a rolling window mean (Fig. 4a, green and red respectively). This revealed divergent complex patterns of neuro-cognitive aging.

**Figure 4:**
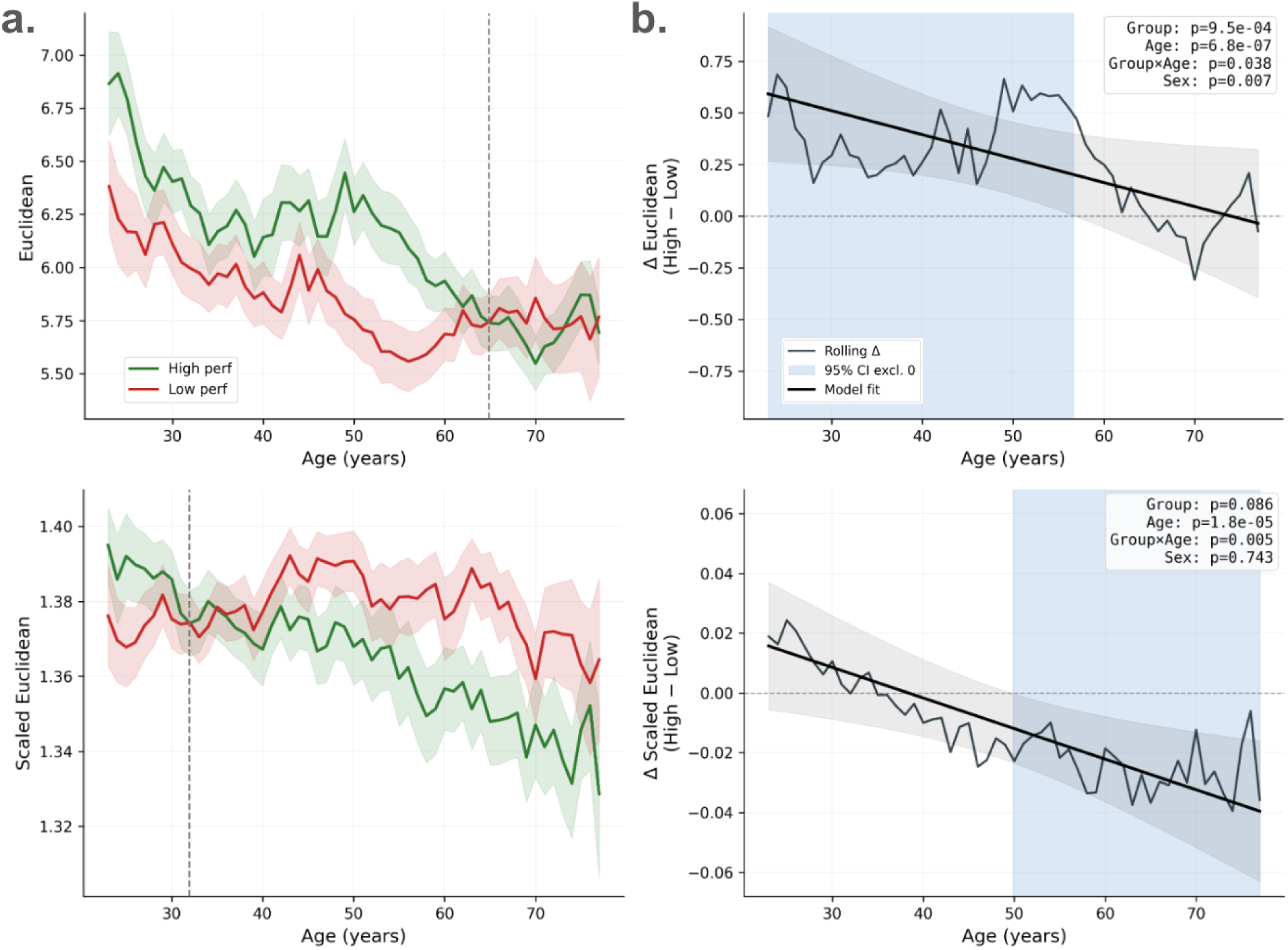
Relationship between neural differentiation, cognition, and age. **a.** Trajectories of average pairwise distance between task-evoked brain activity from 12 task conditions, plotted against age for two distance metrics: Euclidean distance (top), and Scaled Euclidean distance (bottom). The Green curve and shaded area represents the rolling window mean trajectory of high performers (top 50%) and red curve and shaded regions for low performers (bottom 50%), determined by a median split on the age-residualized general cognitive factor. **b.** Rolling window performance group difference (high - low) in differentiation metric (thin black line) with difference model +/− 95% confidence interval overlaid (thick black line and shaded grey area). Light blue shaded region indicates where the 95% confidence interval falls entirely on either side of zero, indicating an age range where the distance metric is significantly different between high and low performers.

For Euclidean distance, high performers maintained elevated neural differentiability across early-to-middle adulthood relative to low performers, with performance groups converging in older age (intersection at 63 years old; Fig 4a, top). This likely reflects a greater dynamic range of activation magnitude across tasks in high performers, which is consistent with greater flexibility, sensitivity to task demands, and adaptability to cognitive workload. However, this advantage diminishes as the capacity for flexible amplitude modulation declines with age (High Performers: r=-0.367, p<0.001; Low Performers: r=-0.191, p=0.019). Importantly, while age-related changes in channel retention and neurovascular integrity may contribute to the overall decline in activation magnitude, they cannot explain the observed differences between high and low performers of the same age.

For Scaled Euclidean distance, the performance groups showed an inverse trajectory throughout aging. In young adulthood, high performers exhibited marginally greater spatial pattern differentiation, but this relationship gradually reversed through middle age (intersection at 36 years; Fig 4a, bottom), such that by older age, high performers showed reduced spatial differentiation across tasks (relative to low performers). In particular, it is the high performers that modulate neural patterns with age, while the low performers exhibit more constant spatial pattern differentiation (High Performers: r=-0.32, p<0.001; Low Performers: r=-0.045, p=0.582). This suggests that in older high performers, maintained cognitive performance is supported by compensatory mechanisms that rely on shared neural scaffolding across cognitive domains. Moreover, preserved or relatively enhanced performance in older age manifests as dedifferentiation in the corresponding patterns of brain activity.

Finally, to formally quantify these observations, we fit an interaction model predicting neural differentiability from Group (High/Low Performance), Age (centered), Group×Age, and Biological Sex as a covariate. We then used the fitted coefficients to derive a difference curve, which was overlaid on the rolling-window difference between high and low performers (Fig. 4b). For Euclidean Distance, the model showed a significant main effect of Age (p<0.001), Group (p<0.001), and Sex (p=0.007). Of particular interest, there was a marginally significant Group×Age interaction (p=0.038), reflecting the aforementioned trend with age for high and low performers to converge. The 95% confidence interval for the difference curve remained above zero until mid 50s (Fig. 4b, top, shaded blue region), indicating that high performers maintained a significant differentiability advantage in the form of a greater dynamic range of task-evoked responses into, but not after, middle age.

Critically, this same period marks a transition in the Scaled Euclidean difference curve (Fig 4b, bottom). As the difference in Euclidean distance (between high and low performers) becomes non-significant, the Scaled Euclidean difference curve crosses into significance in the opposite direction. This is evident as the 95% CI for the Scaled Euclidean distance falls entirely below zero around age 50 and onward. This seemingly synchronized shift suggests that as the capacity to maintain high-magnitude, task-specific activations wanes in late middle age, preserved performance begins to rely on a transition toward compensatory recruitment strategies. Notably, the Scaled Euclidean model is driven by a robust Group×Age interaction term (p=0.005) highlighting the monotonic decrease and eventual sign flip in spatial differentiation between performance groups with age.

We replicated the analysis with deoxygenated-hemoglobin, HbR. While HbR is known to have lower signal-to-noise, it is thought to have better spatial specificity than HbO ^27^. HbR trends mirrored those from HbO (Supplemental Fig. 1 a-c). We also computed reliability of both differentiation scores and found that they are within the typical range of other functional neuroimaging metrics (HbO: Euclidean ICC2=0.503, Scaled Euclidean ICC2=0.293; HbR: Euclidean ICC2=0.561, Scaled Euclidean ICC2=0.321).

Beyond cognitive performance, we repeated this analysis for groups based on subjective markers of mental and cognitive health, measured via PHQ-9, GAD-7, and SCDQ-9 (see Table 1 for summary of scores). Specifically, we categorized participants into ‘None’ (no reported symptoms or cognitive concerns) and Mild+ (mild, moderate or severe symptoms) groups. Since we did not optimize recruitment for target values in these scores we had to rely on natural variance, and thus needed to group all symptoms together to have sufficient power to compare across lifespan (PHQ-9 Mild+ n=90; GAD-7 Mild+ n=110; SCDQ-9 Mild+ n=129).

Repetition of the interaction model for scaled euclidean distance, with these groups as the moderators, revealed evidence that subjective anxiety, depression, and cognitive concerns may alter or interfere with the transition toward compensatory neural mechanisms (Fig. 5a). A consistent pattern emerged; where, individuals with higher clinical scores (Mild+) tended to resist the age-related dedifferentiation seen in the ‘None’ group. While PHQ-9 and GAD-7 scores showed a trend toward a Group×Age (p =0.099, 0.055, respectively), SCDQ-9 yielded a robust, significant interaction (p=0.009) (Fig. 5b). For GAD-7 in particular, the 95% CI fell below zero from middle age onward, suggesting that individuals with higher anxiety maintain higher spatial differentiation, and as they move into older age, fail to engage shared scaffolding observed in the no anxiety group. Likewise, for SCDQ-9 the 95% CI fell below zero, but starting in older age, indicating that recognized cognitive decline was directly reflected in the neural differentiability score of the aged population.

**Figure 5:**
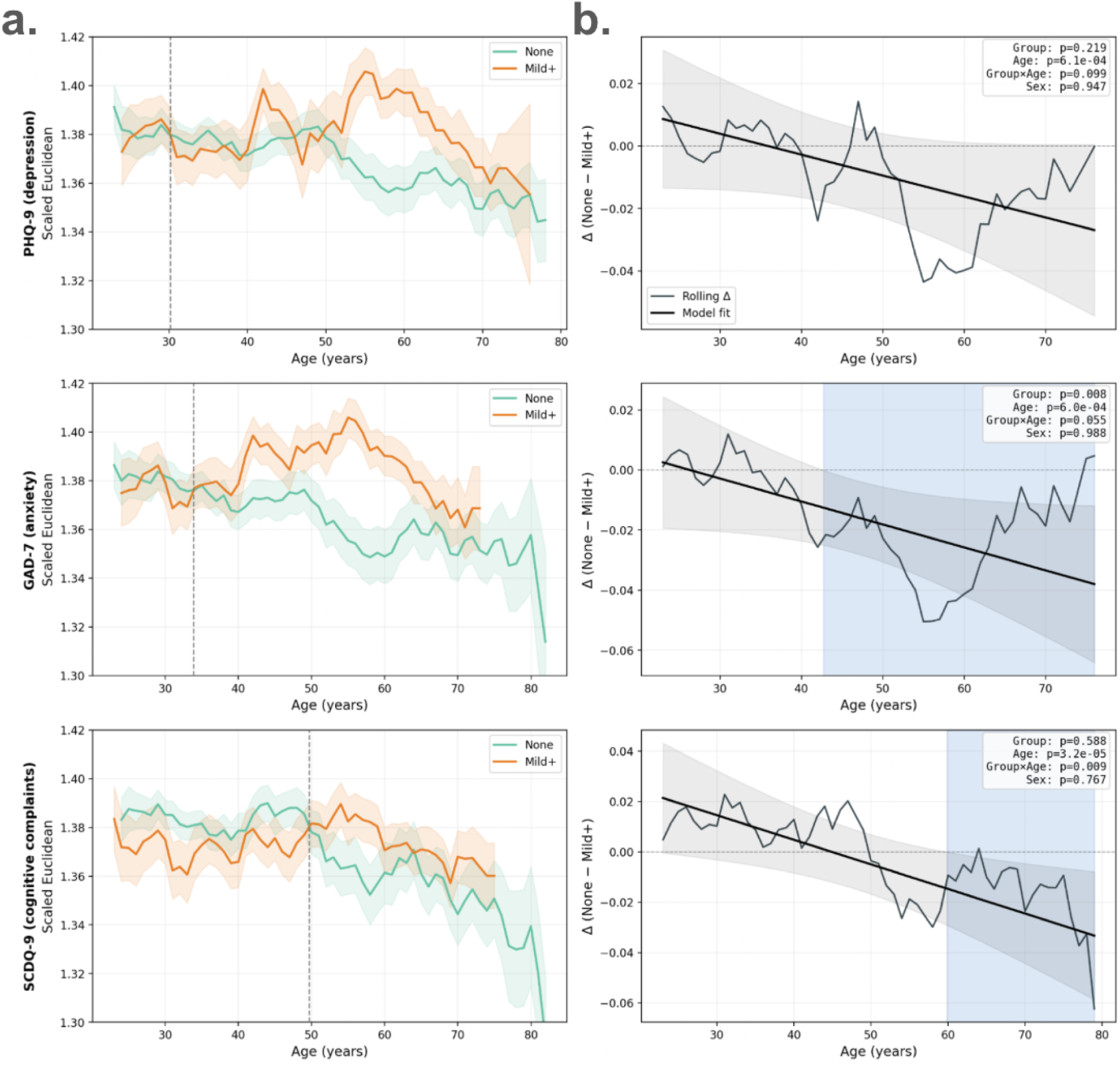
Relationship between neural differentiation, subjective clinical symptoms, and age. a,b. Same as the bottom row of Figure 4, but for PHQ-9 (top), GAD-7 (middle), and SCDQ-9 (bottom). Here teal lines and shaded regions represent no-to-sub-clinical complaints (None) and orange lines and shaded regions represent mild-to-severe complaints (Mild+).

As a control analysis we repeated the subjective complaints group analysis using the GCF (cognitive performance) as the predicted variable (in place of the neural differentiation score). We found no significant main effect of Group or interactions for GAD-7 and SCDQ-9 and only a trend towards a Group×Age interaction (p=0.069) for PHQ-9 that revealed a weak crossover effect where participants with no depression performed slightly worse in younger ages and slightly better in older ages (Supplemental Figure 2). Notably the 95% CI for the model-derived difference curve contained zero across the full age range. The disconnect between our neural and behavioral findings (especially in the case of GAD-7 and SCDQ-9) suggests that either participants’ subjective reports of anxiety or cognitive complaints were reflected in their brain before a measurable decline in cognitive performance, or that the brain is a more sensitive measurement than standard cognitive tasks.

Taken together, these results suggest that magnitude-based and pattern-based neural differentiation follow distinct neuro-cognitive aging trajectories. Namely, while distinct neural circuits and dynamic activations are a hallmark of high cognitive performance into middle age, there is a shift toward recruitment of shared neural circuits in high cognitive performers that persists through older age. Specifically, we have identified a general cognitive pivot in the neural underpinnings of high performance in the early-to-mid 50s, where successful aging becomes characterized by increased spatial dedifferentiation, and that successful aging may be disrupted by depression, anxiety, and other clinical symptoms before behavioral changes can be measured.

## Discussion

In the current study, we have captured changes in cognition with aging using both behavioral and brain-based metrics. We developed a general behavioral measure of cognition across multiple domains that has good reliability over one month. We also developed brain-based metrics showing de-differentiation of brain activity patterns for disparate cognitive tasks with aging. While the behavioral general performance metric showed a gradual and smooth decline with age (Supplemental Figure 3), brain activity followed a more complex pattern separated by performance. High performers showed more variability across task-evoked brain activity until around age 55, suggesting that they have preserved the ability to recruit distinct neural circuits to support specific cognitive tasks, consistent with the maintenance hypothesis ^25^ and CRUNCH^22^. Just prior to this decline, around age 50, high performers start to recruit similar brain networks for different tasks, indicating that these networks may more generally support higher performance at older ages, consistent with theoretical models of neural recruitment in aging, including HAROLD^21^, CRUNCH^22^ and STAC^23^. Taken together, these findings support both the maintenance hypothesis and compensation as critical components underlying cognitive abilities at different time periods in the adult lifespan, with a shift from specialized to general neural populations occurring between early and mid 50s. These findings support the idea that brain metrics provide a more nuanced picture of cognitive function than can be derived from just behavioral metrics.

A number of studies have reported specific ages that mark qualitative changes in brain structure and function. One diffusion tensor imaging study noted major changes in network organization in two phases starting at age 32 and 66^28^.These two age pivot points broadly align with changes we saw between high and low performers, with the intersection point of high and low performers in the Scaled Euclidean age trajectories (age 36) and the convergence of Euclidean age trajectories for high and low performers (age 63). Anatomical pivot points in early 50s^1^ and functional brain changes in the late 40s and early 50s ^5,29^, also align with the pivot period demonstrated in this manuscript, specifically, the end of significant differences in magnitude-based differentiation and the start of significant differences in spatial differentiation between performance groups. Overall the literature broadly supports our findings of change in brain activity patterns that are characterized by phases or pivot points instead of gradual declines.

Extending beyond cognitive performance, we found that subjective reports of anxiety, depression, and cognitive concerns were associated with altered neural aging trajectories. Individuals with higher clinical scores (Mild+) tended to lack the age-related shift toward spatial dedifferentiation observed in high performers. Notably, these group differences and significant interactions with age were apparent in the neural differentiability scores in the absence of significant differences in performance (GCF scores), particularly for GAD-7 (anxiety) and SCDQ-9 (subjective cognitive complaints). This dissociation suggests that neural metrics may be more sensitive than standard behavioral assessments to effects of anxiety and early subjective cognitive concerns. To probe this further, future research should not only selectively recruit participants spanning the full clinical range of depression, anxiety, and severity of cognitive complaints to extend this analysis to more refined clinical grouping (e.g normal, mild, moderate, severe), but also track participants longitudinally.

Strengths of the current study include a large and diverse sample population, the broad multifactor cognitive battery, and the demonstrated reliability of the brain and behavioral metrics. Limitations of this approach include the lack of longitudinal data to link metrics to individual aging trajectories, and the lack of a social cognition task in the cognitive battery.

Overall, the brain-based cognitive battery approach is well-suited for wider adoption and potential routine monitoring of cognitive function. The link between brain metrics and subjective cognitive decline is particularly intriguing, suggesting that these brain-based metrics may be able to objectively measure subjective ratings of cognitive decline and flag patients for early protective interventions. As up to 45% of dementia may be preventable through lifestyle interventions ^30^, identifying the patients most likely to benefit could appreciably reduce dementia incidence. Future work following patients longitudinally and linking the metrics to clinical conditions beyond healthy aging (i.e. mild cognitive impairment) would provide a foundation for a future where brain health can be extended due to precision early intervention.

## Methods

### Participants and screening procedures

Healthy participants (N=302, 141 female, mean age=48.2, range 18-87) completed one study visit. A subset of participants (n=106, 50 female, age=44.6 ± 15.2, range 18-74) completed a second study visit roughly one month later (time between visits: mean±STD=23.1 ± 8.3 days, range=8–43 days).

Participants were screened using the following inclusion criteria: (1) at least 18 years or older at the time of enrollment, (2) ability to perform informed consent on their own, (3) fluency in spoken and written English, (4) willingness to allow anonymized photos of the face and head taken during some study procedures to verify headset placement. Additionally, participants were screened out based on the following exclusion criteria: (1) prior diagnosis of mild cognitive impairment (MCI) or other memory impairment, (2) first-degree relative with dementia or clinically relevant memory problems; (3) Alzheimer’s or dementia diagnosis, (4) uncorrected major visual or auditory deficits that would prevent them from completing a study task, (5) current or recent (in the past 6 months) chemotherapy and/or radiation for any cancer, and (6) major medical or psychiatric conditions associated with secondary causes of cognitive decline, including but not limited to uncontrolled diabetes, hypertension, or hypothyroidism; Parkinson’s disease; motor neuron disease; multiple sclerosis; brain tumor; stroke; encephalitis; meningitis; epilepsy; traumatic brain injury with severe sequelae (e.g., coma, loss of consciousness >2 hours, or skull fracture); or current severe psychiatric disorders (e.g., bipolar disorder, schizophrenia, or psychosis).

Participants gave written informed consent before beginning the study. The study was reviewed for ethics by the Advarra IRB, which approved this study (#Pro00083567). All experimental protocols were approved by the IRB and were conducted in accordance with the Declaration of Helsinki. Participants received monetary compensation for their time, effort, and travel expenses.

Five additional participants were recorded that are not included in the analysis. Reasons for exclusion include: one participant who experienced headset discomfort, one participant with an illness (coughing fits during recording), two participants with two hardware issues during recording, and one participant with no responses recorded during the last two of the tasks.

### Cognitive multifactor task design

The task (Fig. 6), developed and presented using the Unity game engine, consisted of six subtasks administered sequentially as a self-paced “monotask.” Each subtask began with on-screen instructions or a brief guided tutorial, which participants completed at their own pace. Participants were allowed to take breaks between subtasks and, in some cases, between blocks within a subtask, resulting in variation in total task duration across individuals (mean ± STD = 28.1 ± 1.8, range 24-39 minutes).

**Figure 6:**
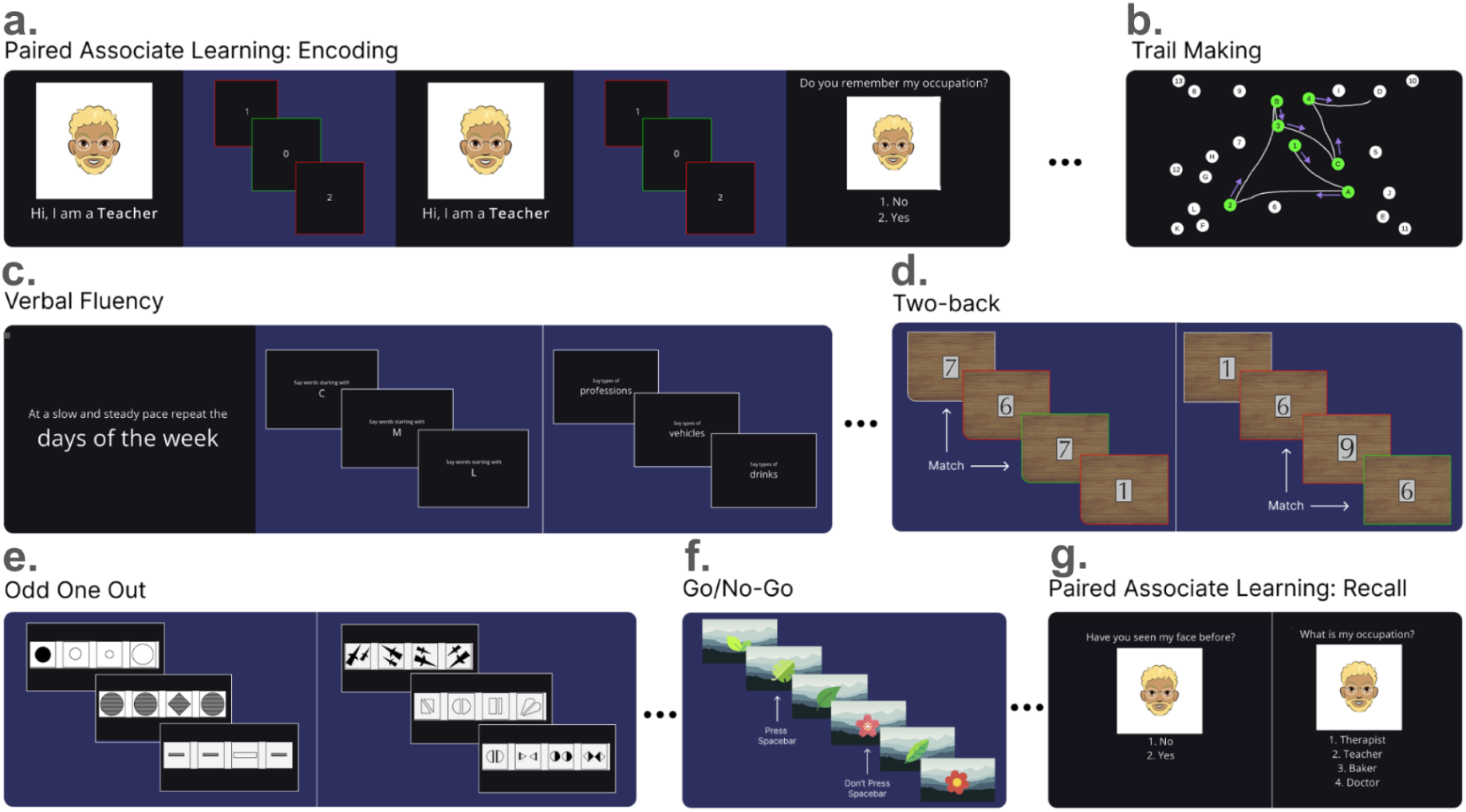
Multifactor task design. **a.** Paired Associate Learning task, encoding phase. **b**. Trail making task. **c.** Verbal Fluency task. **d**. Two-back task. **e**. Odd One Out task. **f.** Go/No-Go task. **g**. Paired Associate Learning task, recall phase.

Before the start of each subtask, participants were instructed to close their eyes for 20 seconds and then open their eyes for 20 seconds. The subtasks are described below in the order of presentation.

#### Paired Associate Learning (PAL) – Encoding

The PAL task (Fig. 6a) consisted of two components, one at the beginning of the session and one at the end.

In the initial PAL Encoding phase, participants viewed pairs of items and were asked to memorize the association between them. Each pair consisted of a cartoon person and that person’s occupation. Twelve pairs were presented twice, with each presentation lasting 4 seconds.

After each encoding round, participants completed a distractor task in which single-digit numbers appeared in rapid succession. Participants pressed the spacebar whenever a 0 appeared. Each distractor block consisted of 40 trials, with stimuli presented for 250 ms and a 500 ms inter-trial interval (ITI). The total task took approximately 2 minutes 45 seconds.

#### Trail Making

The Trail Making Test (Fig. 6b) was a computerized adaptation of the standard paper-and-pencil version (TMT-B). Participants viewed 26 circles displayed on the screen, each containing either a number (1–13) or a letter (A–K). They were instructed to move the mouse to connect the circles in alternating numeric and alphabetic order (1–A–2–B–3–C, etc.).

If participants passed through a circle out of sequence, an audible buzzer sounded and an on-screen correction prompt appeared. Participants were given 2 minutes 30 seconds to complete the task, but the task was considered complete if the sequence was properly traced before that.

#### Verbal Fluency

In the Verbal Fluency task (Fig. 6c), participants generated as many words as possible in response to a prompt on the screen. The control condition required participants to repeatedly recite the days of the week. The experimental conditions asked participants to say words starting with a specified letter (phonological prompts) or belonging to a semantic category.

The task began with the 20-second control condition, followed by three phonological prompts (20 seconds each) and three semantic categories (20 seconds each). Participants pressed the spacebar each time they produced a word. Because speech was not recorded, this provided a rough count of verbal output. The task took approximately 2 minutes 20 seconds to complete.

#### Two-Back

This task was a modified version of the N-back paradigm (Fig. 6d). Numbered playing cards appeared sequentially on the screen, and participants responded by pressing the spacebar whenever the current card matched the card presented two positions earlier.

The task consisted of 4 blocks, each containing 26 trials. Each card was displayed for 700 ms, followed by a 500 ms ITI. Each block contained 25% target (match) trials. The task took approximately 2 min 15 seconds to complete.

#### Odd One Out

In this task (Fig. 6e), participants viewed sets of four images and selected the one that differed from the others. The task included two “easy” blocks and two “hard” blocks. Participants were instructed to complete as many trials as possible within the time allotted for each block, which was 30 seconds. The task took approximately 2 minutes 15 seconds to complete.

#### Go/No-Go

In the Go/No-go task (Fig. 6f) participants viewed a series of images (leaves and flowers) presented sequentially. They pressed the spacebar when a leaf appeared (go trials) and were instructed to press nothing when a red flower appeared (no-go). Feedback was provided via tones: a pleasant tone for correct responses (hits) and an unpleasant tone for false alarms.

The task consisted of 4 blocks, each with 40 trials. Stimuli were displayed for 400 ms, followed by an ITI of 600 ms ± 100 ms. Participants were required to respond within the 400-ms presentation window. No-go trials (flowers) constituted 25% of each block. The task took approximately 3 minutes to complete.

#### PAL – Recall

As the final subtask, participants completed the PAL Recall phase (Fig. 6g), assessing memory for associations learned during PAL Encoding. Two conditions were administered: recognition (new/old) and forced-choice cued recall. In the recognition condition, participants were shown 24 faces (12 previously seen, 12 novel) and indicated whether each face had appeared during encoding. In forced-choice cued recall, participants viewed the 12 original faces, each paired with four occupation options (one correct), and selected the occupation originally associated with each face.

Participants had up to 8 seconds to respond on each trial. The ITI was 500 ms, measured from the time of response. Task time was variable, with a maximum total task time of 4 minutes 48 seconds.

#### Behavioral Quantification and Synthesis

For each task condition, Principal Component Analysis (PCA) was run on the full set of behavioral metrics, reducing the data to a small number of components (typically 1–3) that captured the primary dimensions of task performance. Parallel Analysis was employed to determine the number of PCs to retain for each task condition. These PCs were inputs to a second-level PCA, and the resulting first PC, termed the General Cognitive Factor (GCF), was used in subsequent analyses.

### fNIRS data collection and feature extraction

#### Quantification of Task-evoked Brain Activity

TD-fNIRS measurements were performed with the Kernel Flow system^26^. The headset has a modular design, such that each module unit contains 3 light sources, 6 detectors, and covers a discrete scalp region. Together the headset records data over the whole head including the prefrontal, parietal, temporal and occipital cortices. Data preprocessing including bad channel removal, conversion to Hb values, motion correction, detrending, and short channel regression, followed previously published methods^16,31–33^.

Task-evoked hemodynamic activity was quantified using generalized linear models (GLMs) for each task condition as described in Dubois et al. (2024, 2025). In total, 12 task conditions were included in the design matrices, namely: go-no-go, 2-back, encoding, distractor, forced choice cued recall, recognition (new/old), semantic, phonological, control (fluency repetition), tmt-B, easy odd-one-out, and hard odd-one-out. Group-level activations were computed using standard statistical tests (two-sided independent t-tests) on individual GLM beta weights. For individual and group-level analyses, the resulting test statistics for channels with source-detector-separation of > 15 mm, were pooled according to the module the channel-source belonged to, yielding 70 measurements (35 regions x 2 Hbs) for each task condition. These regional activations were used for all further analyses.

#### Neural Differentiation Metrics

For each participant, task-evoked brain activity (for HbO and HbR separately) was represented as a vector in a 35-dimensional space, where each dimension corresponds to a headset module (brain region as described above), and pairwise Euclidean distances between all task condition vectors were computed. A general neural differentiation score was derived for each participant as the average (mean) distance across all task condition pairs. A second metric, Scaled Euclidean distance, was computed identically but each participant’s task condition vectors were first divided by the mean vector norm across all task conditions within that session. This normalization removed session-level differences in overall activation magnitude, isolating the contribution of spatial activation patterns to neural differentiation. As with Euclidean distance, a general Scaled Euclidean differentiation score was then calculated as the average scaled distance across all task pairs. Note that for both metrics, lower values indicate more similar (less differentiated) neural representations across tasks.

#### Evaluation of Age and Performance on Neural Differentiation

Age-related trends and performance-related trends in neural differentiation scores were assessed using ordinary least-squares regression (OLS model). Specifically, each distance metric was modelled as a function of age (mean centered), performance group (High vs Low determined by a median split on age-detrended GCF scores), their interaction, and biological sex included as a covariate. P-values of fitted coefficients were used to evaluate the significance of main effects and whether age-related change in neural differentiation differed across performance groups (the interaction term). From the fitted model we derived a High – Low difference curve and used the 95% confidence interval of the difference curve to identify age ranges where neural differentiation significantly differed between performance groups. We repeated this analysis for subjective reports of depression (PHQ-9), anxiety (GAD-7), and cognition (SCDQ-9), where groups were defined as None (no complaints) and Mild+ (mild-to-severe complaints).

## Data Availability

The data that support the findings of this study are not openly available due to reasons of sensitivity and are available from the corresponding author upon reasonable request and with a data sharing agreement.

## Code Availability

The underlying code for this study is available to approved researchers after completing a Material Transfer Agreement.

## Acknowledgments

We thank the participants for their time and contribution to our study. This study was funded by Kernel.

## Author Contributions

Conceptualization: KLP, RMF. Software: JD, EMK, ZMA, GL. Methodology: JD, EMK, ZMA, KLP. Formal analysis: EMK, JD. Investigation: YC, DD, AG, AJ, EMK, NM, MT. Resources: YC, DD. Project Administration: MT, KLP. Data curation: NM. Visualization: EMK, NM. Writing—Original Draft: EMK, KLP, NM, MT. Writing—Review and Editing: all authors. Author order was determined alphabetically. Dr. Katherine Perdue agrees to be accountable for all aspects of the work, ensuring that questions related to the accuracy or integrity of any part of the work are appropriately investigated and resolved.

## Supplemental figures

**Supplemental Figure 1:**
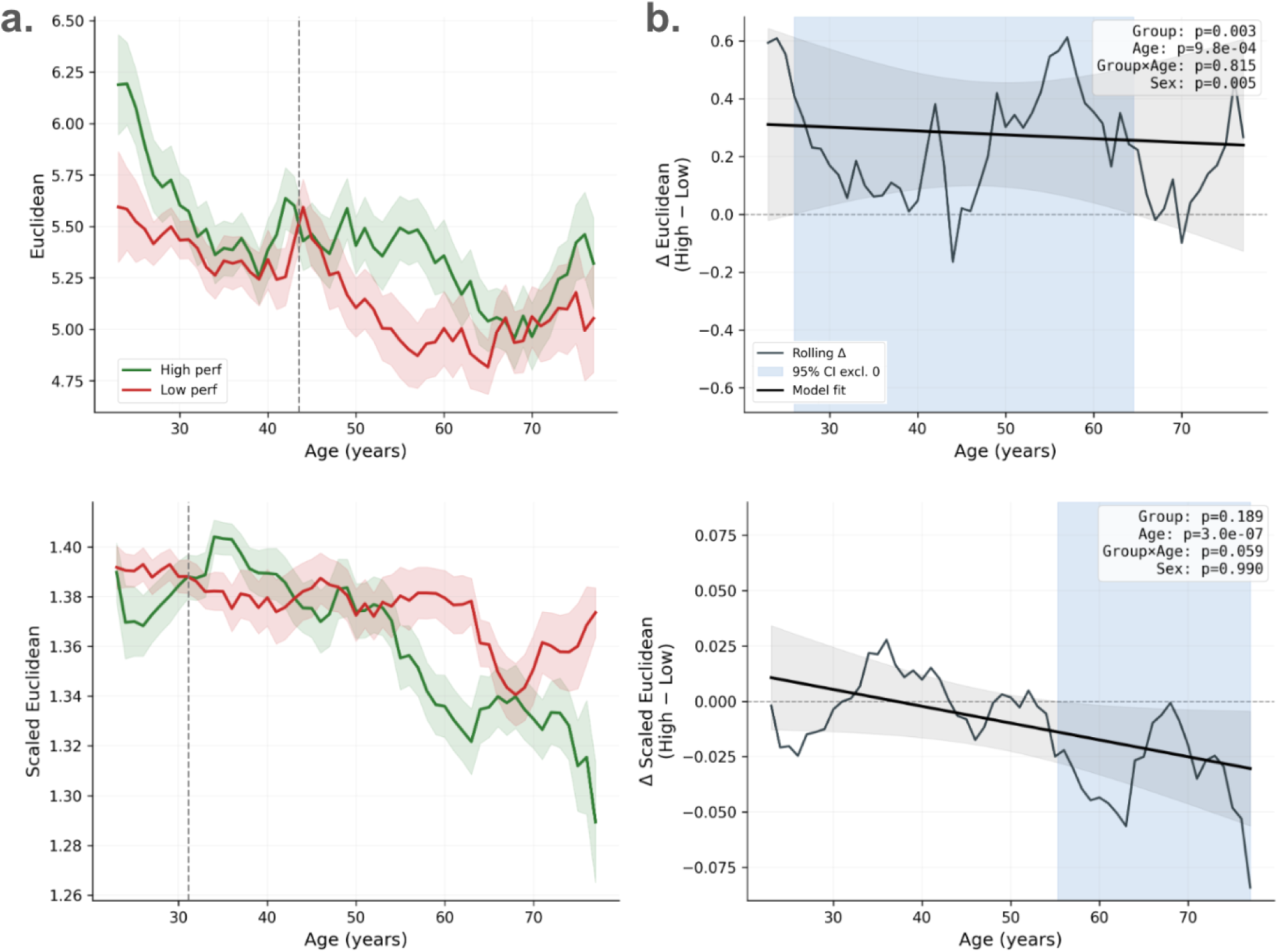
Replication of differentiation results using HbR. Same as main Figure 4, but for HbR.

**Supplemental Figure 2:**
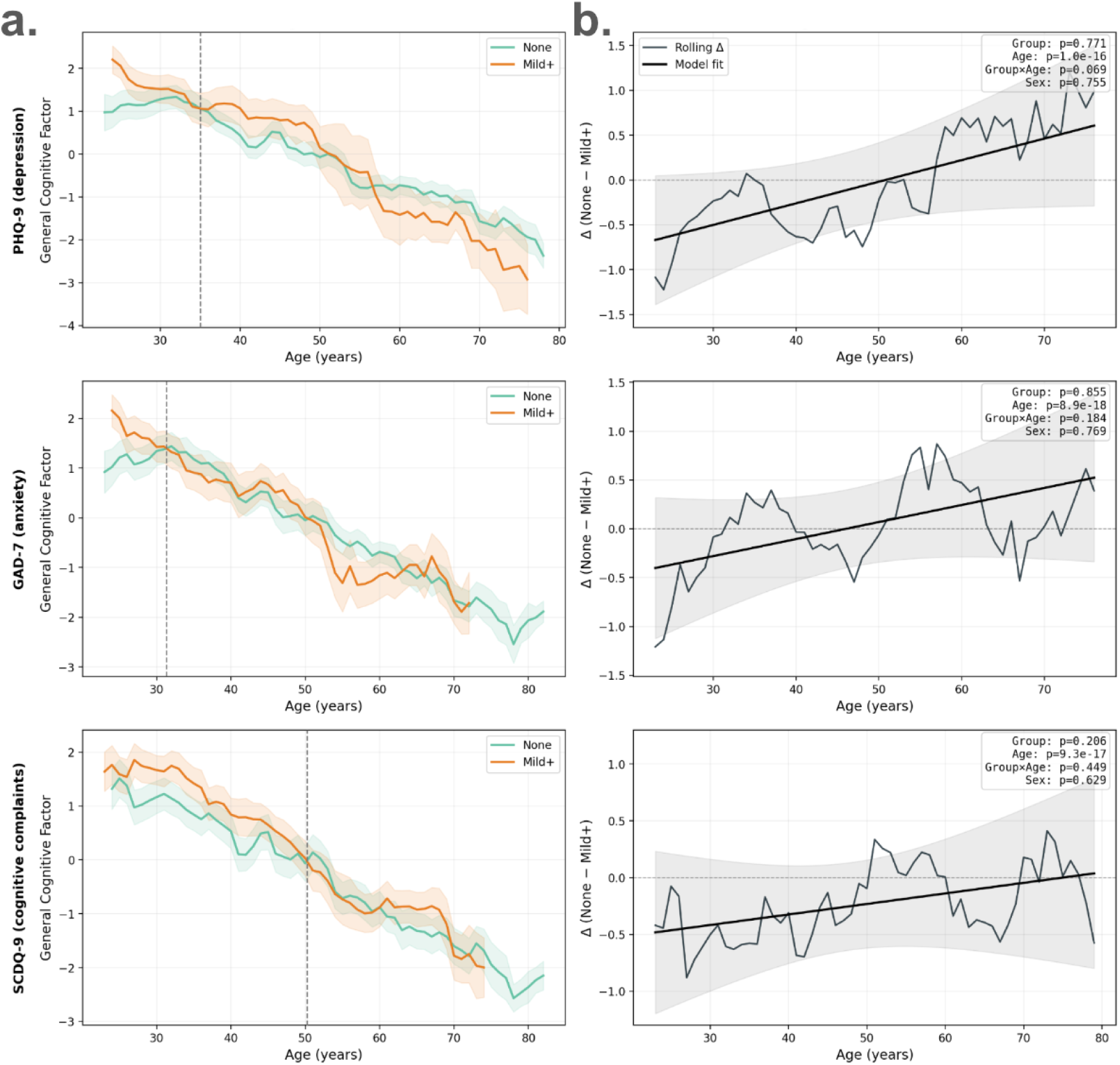
Same as main Figure 5, but with the GCF (General Cognitive Factor) as the predicted variable. Note that the only significant effect across all surveys is the main effect of age.

**Supplemental Figure 3:**
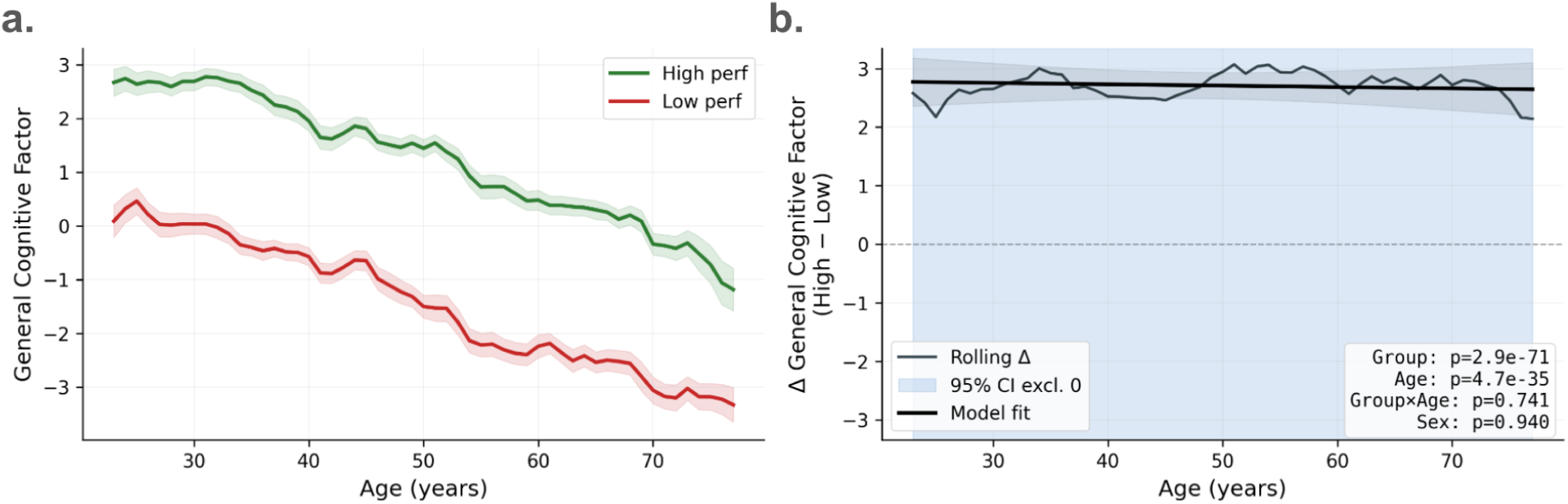
What does it mean to be a high performer within each decade? **a.** Rolling window average GCF (General cognitive factor)) plotted against age, separated by age-residualized performance group. High performers are shown in green (top 50%) and low performers in red (bottom 50%). Note the consistent relationship with age, with both groups declining at a similar rate. Despite the trend with age, the large separation between performance groups demonstrates that a high performer at age 60 may have better performance than a lower performer below the age of 30. **b.** High performers less low performers difference plot (thin black line) vs age. A difference curve derived from an interaction model is overlaid (thick black line). The gray shaded region represents the 95% CI. The blue shaded band represents the age range for which the difference (between high and low performers) is significant (does not contain 0). Note that the difference is flat, and while there are robust main effects of Group and Age the interaction term is not significant. This is also reflected in the flat (constant) nature of the fitted difference model.

