## Supplemental Figures for "Brain dynamics supporting high cognitive performance reorganize after midlife"

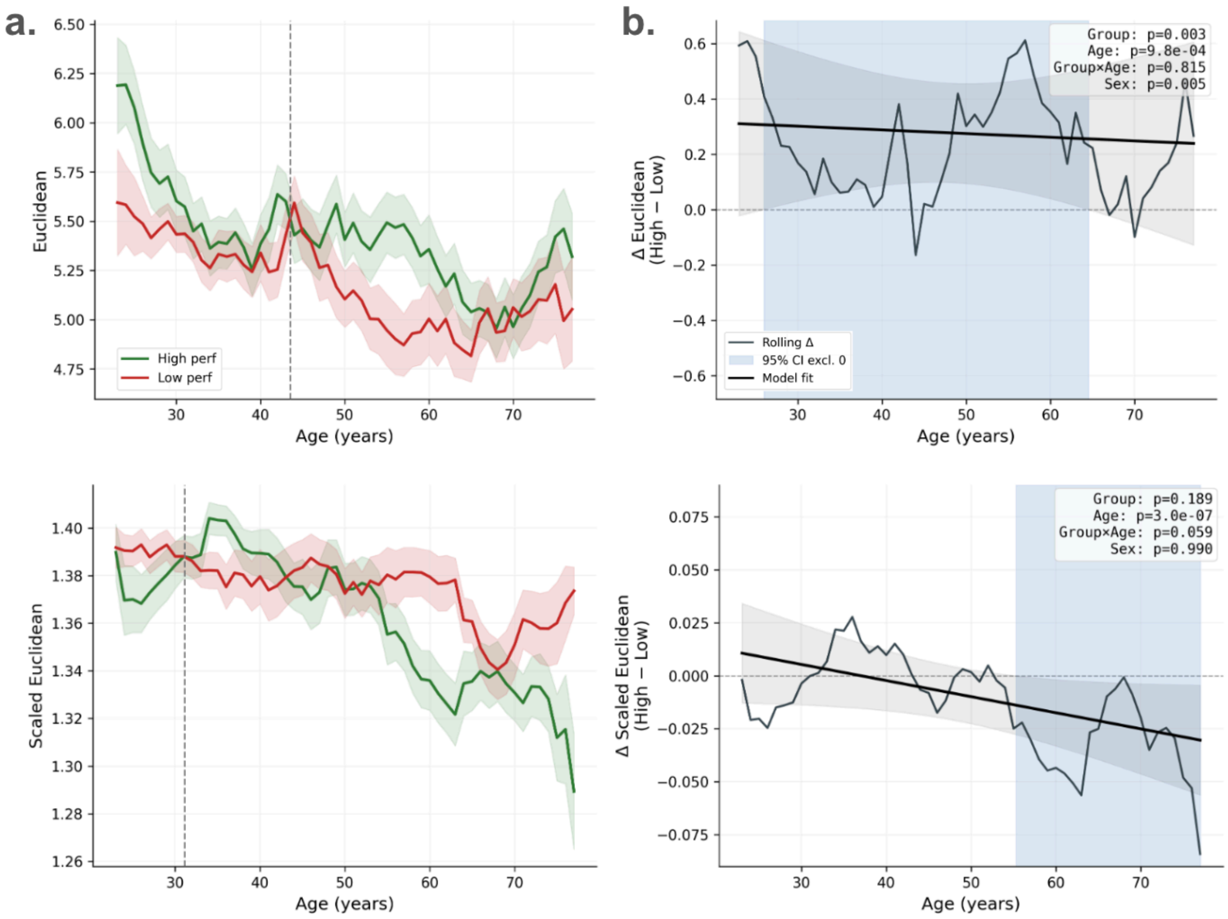


**Supplemental Figure 1: Replication of differentiation results  using HbR.**
Same as main Figure 4, but for HbR.

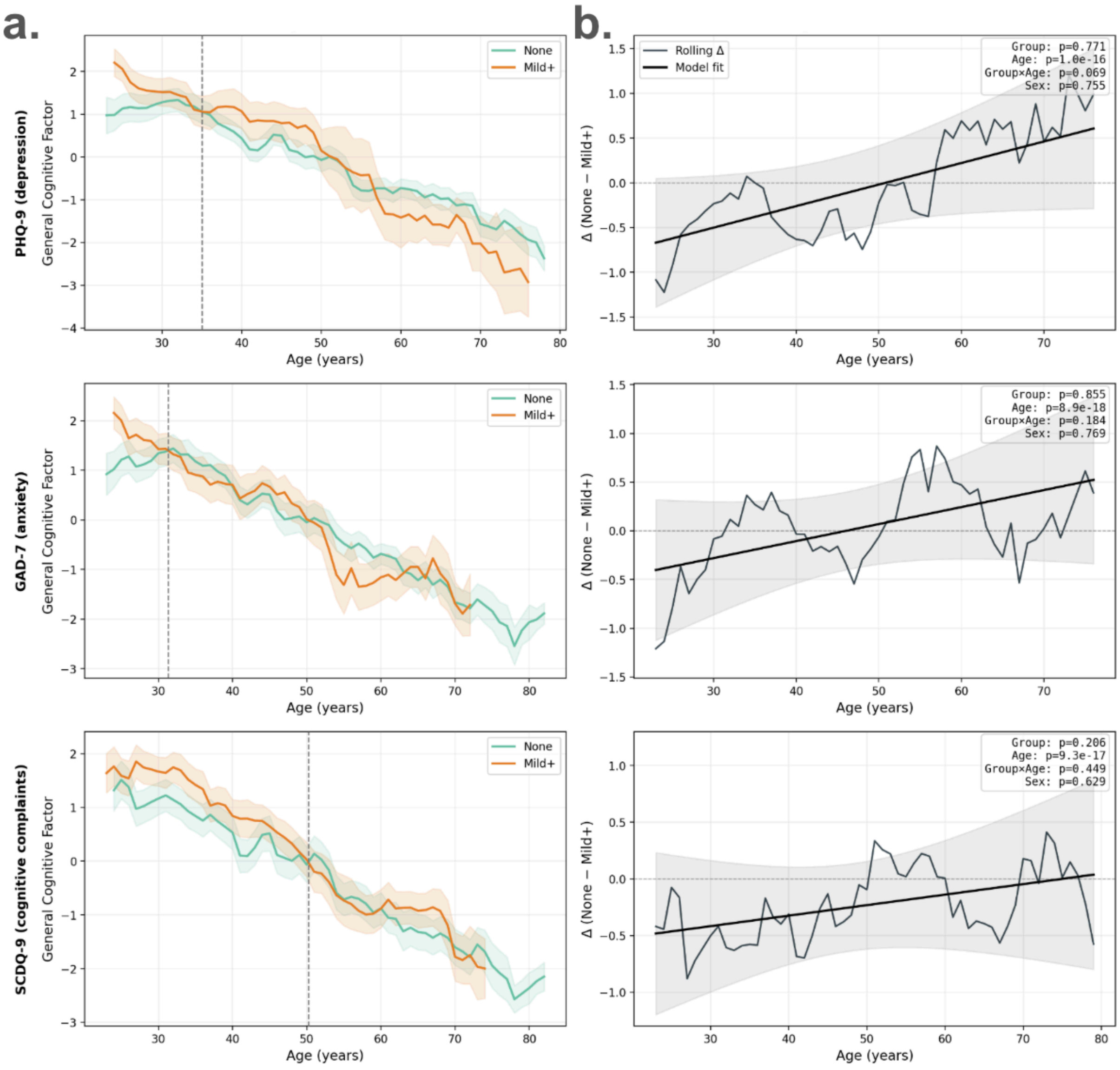


**Supplemental Figure 2:**
Same as main Figure 5, but with the GCF (General Cognitive Factor) as the predicted variable. Note that the only significant effect across all surveys is the main effect of age.

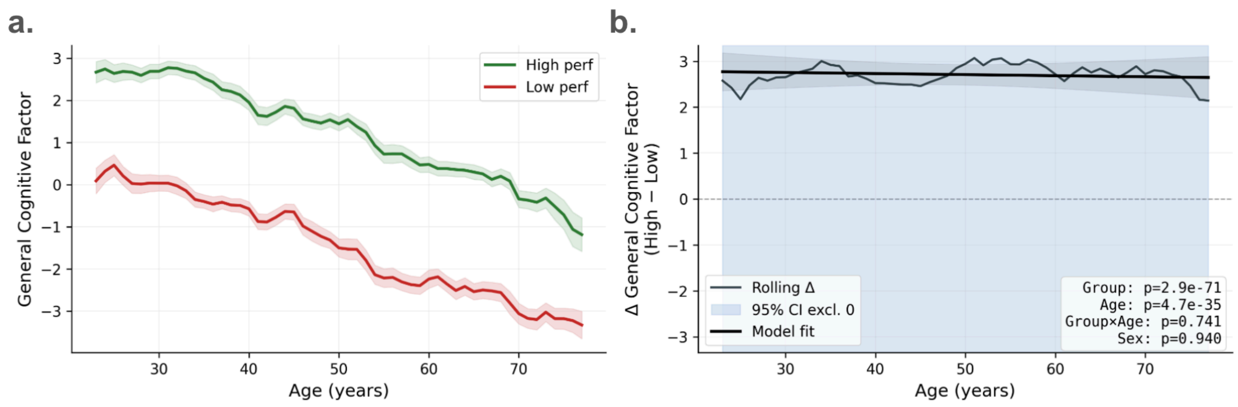


**Supplemental Figure 3: What does it mean to be a high performer within each decade?**

**a.** Rolling window average GCF **(**General cognitive factor)) plotted against age, separated by age-residualized performance group. High performers are shown in green (top 50%) and low performers in red (bottom 50%). Note the consistent relationship with age, with both groups declining at a similar rate. Despite the trend with age, the large separation between performance groups demonstrates that a high performer at age 60 may have better performance than a lower performer below the age of 30.

**b.** High performers less low performers difference plot (thin black line) vs age. A difference curve derived from an interaction model is overlaid (thick black line). The gray shaded region represents the 95% CI. The blue shaded band represents the age range for which the difference (between high and low performers) is significant (does not contain 0). Note that the difference is flat, and while there are robust main effects of Group and Age the interaction term is not significant. This is also reflected in the flat (constant) nature of the fitted difference model.
